# The Importance of Reverse Translation for Preclinical Off-Target Mitigation: Quantification and Mitigation of Biases in the FDA Adverse Event Reporting System

**DOI:** 10.1101/068692

**Authors:** Mateusz Maciejewski, Eugen Lounkine, Steven Whitebread, Pierre Farmer, Bill DuMouchel, Brian K. Shoichet, Laszlo Urban

**Author notes:** Corresponding authors: Laszlo Urban, MD, PhD, Preclinical Secondary Pharmacology, Novartis Institutes for BioMedical Research, 250 Massachusetts Avenue, Cambridge, MA., 02139, U.S.A.; Mateusz Maciejewski, PhD, Inflammation & Immunology, Pfizer Inc., 1 Portland Street, Cambridge, MA, 02139, U.S.A.; Eugen Lounkine, PhD, Center for Proteomic Chemistry, 250 Massachusetts Avenue, Cambridge, MA., 02139, U.S.A.; Brian K. Shoichet, PhD, Dept. of Pharmaceutical Chemistry, University of California, San Francisco, 1700 4th St., Byers Hall Suite 508D CA., 94158-2550, U.S.A.

## Abstract

The Food and Drug Administration Adverse Event Reporting System (FAERS) is the primary source for post-marketing pharmacovigilance. Though potentially highly useful, the database reflects reporting biases, stimulated reporting, and suffers from lack of standardization and the use of multiple drug synonyms. These biases can suggest adverse drug reactions (ADRs) where none exist, and can obscure others that do exist. To decrease the noise in FAERS, and to reinforce important associations, we mapped over 750,000 drug identifiers in FAERS to the normalized chemical structures of their ingredients. This illuminated associations that would not otherwise be apparent, and also allowed a time-resolved analysis of ADR reporting. It also revealed similarities between drugs and adverse events across therapeutic classes, enabling unbiased classification of adverse events, indications, and drugs with similar clinical profiles. For instance, comparison of two selective cyclooxygenase-2 inhibitors, celecoxib and rofecoxib finds distinctive FAERS profiles after time-resolved analysis. We also investigated key idiosyncrasies, such as confusion between drug indications and drug ADRs, which can tar a drug treating a life-threatening disease, like thalidomide’s use against myeloma, with a deadly ADR that is likely the result of the disease itself, multiplications of the same report, which unjustifiably increases its apparent importance, and the correlation of reported ADRs with public events, regulatory announcements, and with publications. Comparing the pharmacological, pharmacokinetic, and clinical ADR profiles of methylphenidate, aripiprazole and risperidone, and of kinase drugs targeting the VEGF receptor (VEGF-R2), demonstrates how underlying molecular mechanisms can emerge from ADR co-analysis. The precautions and methods we describe may enable investigators to avoid confounding chemistry-based associations and reporting biases in FAERS, and illustrate how comparative analysis of ADRs can reveal underlaying mechanisms.

## INTRODUCTION

Safety assessment of drug candidates is crucial for drug discovery, enabling the development of medicines that achieve the desired therapeutic effects with the least risk of adverse side effects. Preclinical regulatory investigations and clinical trials are designed to address safety of drug candidates and eliminate those that do not meet risk-benefit expectations^1^. However, limited access to large, diverse patient populations in clinical trials, untested drug co-administrations, and development of ADRs associated with chronic treatment, often results in post-marketing labeling and occasional withdrawals^2–4^. Thus, postmarketing pharmacovigilance is essential to track ADRs and ultimately reduce the over 1 million serious drug related side effects that occur each year in the USA. Between 5-10% of these ADRs are fatal^5^, and many others cause patient suffering, hospitalization, and increased health system burden^6^. Indeed, the fatality rate attributed to ADRs puts them among the top causes of death in the USA (over 40,000 in 2011), similar to suicide-related mortality^7^.

Determinant tools in post-marketing pharmacovigilance are databases that aggregate ADR reports. Key among these is the FDA Adverse Event Reporting System (FAERS), which is perhaps the most extensive, and among the most widely accessible of these databases, currently containing over 8.5 million reports and rapidly growing^8^. FAERS and related databases, such as those of the EMEA and of Health Canada, can provide specific ADR phenotypes typical for either individual drug classes or specific indications, and can be accessed either directly^8^ or by APIs ^9,10^. These large-scale adverse event databases enable analysis to relate clinical phenotypes and compounds^11^, and they have been widely used by the clinical community with much impact^12–15^.

It is an attractive proposition to exploit the sheer scale of FAERS to detect drug-ADR associations that would otherwise not be apparent. A challenge in doing so has been the heterogeneous data sources and data conflation in the database.

FAERS, while providing a solid frame for reporting, contains redundancies, biases, and conflations that affect its analysis and interpretation^16^. Our ability to even correlate drugs with their effects is obscured by something as simple as the tangle of drug synonyms in FAERS - on average 16 different names for medicines containing each active drug ingredient - which can obscure associations.

In this work, we investigate the effects that these data conflations, inflations, and inaccuracies can have on ADR and mechanistic inference from FAERS, and methods to address them. We begin by mapping drug identifiers in FAERS to normalized chemical structures of their ingredients, which brings together observations over the “full drug”, not just particular drug names and synonyms, which remain incomplete. With chemical structure analysis in hand, we were able to compute time-resolved profiles of drug-ADR associations, which revealed intriguing comorbidities and similarities of ADRs between drugs, and of their time evolution. We then turn to the origins and control for reporting biases in FAERS, considering stimulated reporting and the several different, often non-medical communities that can contribute to FAERS, something itself facilitated by the time-evolution analysis of ADR reports, and its correlation with contemporary news events. We illustrate how these biases can ramify with in depth analysis of FAERS content on two COX-2 inhibitors, rofecoxib and celecoxib, and with two PPAR-γ agonists, rosiglitazone and pioglitazone. As examples of how these analyses can link ADRs to specific targets, we consider the differential ADR profiles of drugs used for the treatment of the Attention Deficit Hyperactivity Disorder (ADHD), and their distinct ADRs may be explained partly by molecular targets - a logic that is often used - combined with pharmacokinetic exposure - which is often overlooked. Similarly we investigate the differentiation of the hypertensive side effects of VEGF-R2 inhibitors based on their potency and pharmacokinetic (PK) profile. The precautions and methods we describe, may enable investigators to use FAERS at increased confidence and avoid confounding chemistry-based associations and reporting biases in FAERS, and illustrates how comparative analysis of ADRs can reveal underlying mechanisms and highlight the reverse translation value in the drug discovery process.

## RESULTS

### Analysis of content: Unexpected trends in FAERS reporting

The FAERS database holds over 8.5 million reports and is steadily growing (over 1,320,000 reports added in 2015; Figure 1A. We extracted 8,749,375 FAERS reports, mapped to 7,095,566 individual cases. Often a patient’s condition is monitored over a span of multiple reports, which must be considered when investigating the incidence of a particular drug-ADR association^8^.

**Figure 1.**
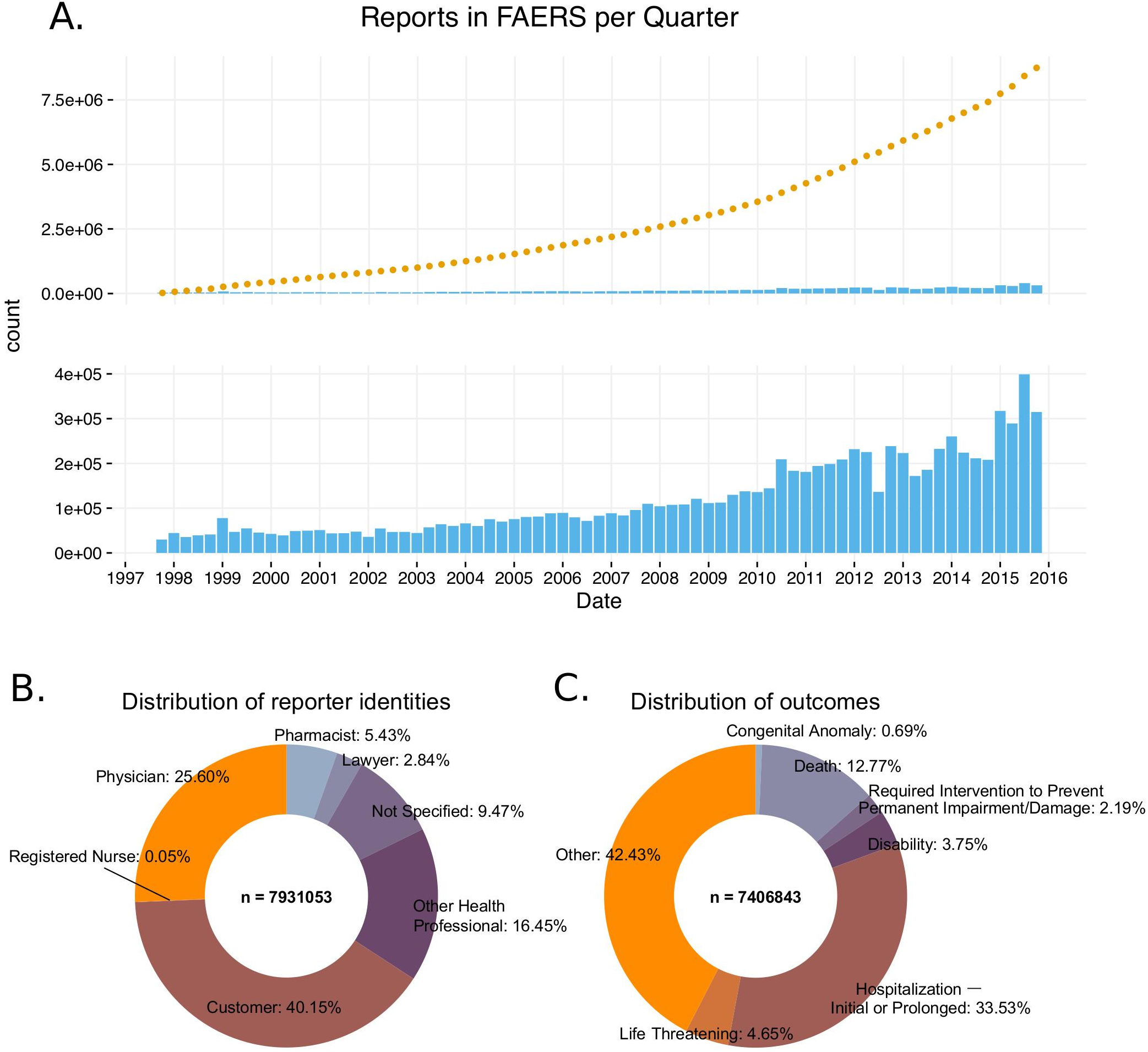
General information of the FDA Adverse Event Reporting System (FAERS) content (1997-2015). A) The cumulative number of reports in FAERS is shown in the top panel; the bottom panel shows the number of new reports per quarter. B) Distribution of reporter identities. Data are based on reports submitted between Q2 2002 (identification of reporting individuals started at this time), and Q4 2015. C) Distribution of reports by the 7 ADR outcomes defined in FAERS.

Inflation of reports by multiplication can increase the apparent significance of a drug – adverse effect association, particularly when the total number of reports is low. To systematically identify the most similar cases, we compared all pairs of reports using demographic and prescription data. Almost 1% of the reports in FAERS (61,780 cases) represent multiple entry cases with identical drugs, identical ADRs, event dates, patient age and gender (Supplementary File 1). Intriguingly, only half of the reports were submitted by health-care professionals (Figure 1B). Over one-third (3.2 million) were initiated by the patients themselves and 9% were labeled “non specified”. Lawyers reported 3% of all FAERS cases (Figure 1B).

FAERS uses seven descriptors of report outcomes: “Death”, “Life-Threatening”, “Disability”, “Congenital Anomaly”, “Required Intervention to Prevent Permanent Impairment/Damage”, “Hospitalization – Initial or Prolonged”, and “Other”. Among these, only “Other” is used to report relatively benign outcomes. Unexpectedly, such benign outcomes are reported only around 40% of the time, whereas almost 15% of reported cases result in death (Figure 1C). It is a feature of reporting in an open submission database like FAERS that this ratio does not reflect the true balance between fatal and relatively benign drug ADRs, but rather the ratio of the ADRs that are thought to merit reporting.

Among the 945,526 reports where death is the outcome of the ADR, 42,526 were linked to cardiac arrest, and 50,155 to suicide. Top molecular ingredients of drugs that were primary suspects in death reports were rosiglitazone: 17,165 (indication type II diabetes), rofecoxib: 11,386 (primary indications: arthritis, pain; withdrawn from the clinic), reteplase: 11,386 (indication of acute myocardial infarction (MI)), and thalidomide: 17,104 (indication of myeloma multiplex; additionally, 26,429 cases of death have been attributed to lenalidomide, a derivative of thalidomide also prescribed for myeloma). For drugs like rofecoxib or rosiglitazone, which are prescribed for manageable and non-life threatening diseases, the inference that the ADR has lead to death can be reasonably made. Similarly, a comparison of celecoxib (reported number of deaths: 4,066; Standardized Mortality Ratio [SMR]^17^: 1.3) and rofecoxib, which are prescribed for the same indication, highlights the significantly higher SMR of patients taking the latter drug (SMR: 5)^18^. However, the attribution of death as an ADR of thalidomide when it is used to treat myeloma multiplex, a life threatening, malignant disease^19^ may be hard to support; it seems likely that the “ADR” here reflects the cancer that the drug is meant to treat. Similarly, the acute myocardial infarction that reteplase is used to treat^20^ may well be the cause of many of the death ADRs with which the drug is tarred, not the drug itself. When a drug is used to treat a life-threatening disease, care is warranted in interpreting death as an ADR of that drug.

### Mapping drugs to their molecular ingredients improves signal retrieval

In most FAERS studies, drugs are identified using R_x_Norm^21,22^, a set of drug synonyms supplied by the National Library of Medicine. This mapping is sufficient for the questions that may be asked of FAERS by a clinical professional, such as the safety signals for a particular drug formulation. However, products that have different identities in resources such as R_x_Norm share common molecular ingredients and are highly similar in their effect on molecular targets. To investigate the ADRs associated with fluoxetine, for instance, one must aggregate its 378 different synonyms. Without such aggregation, well-known fluoxetine side effects such as sexual dysfunction become statistically insignificant (4 cases when only the fluoxetine drug synonym Prozac is considered; Relative Reporting Ratio [RRR] = 1.75; q-value = 1), whereas once aggregated, these ADRs stand out clearly (87 cases; RRR = 6.67; q-value = 2.56·10^-96^). Conversely, in its non-aggregated form, Prozac appears to have statistical significant associations with sex chromosome abnormality (1 case; RRR = 2.96; q-value = 2·10^-3^). Aggregated, however, this association becomes insignificant (1 case; RRR = 2.78; q-value = 1). For those interested in the molecular bases of drug actions and side effects, a simple way to interrogate the drugs as molecules is critical.

Accordingly, we mapped the active drug ingredients in over 98% of the reports using a combination of natural language processing and multiple databases of synonyms (see Methods). Not only does this value compare favorably to the 81% recognition achieved using only the synonyms in R_x_Norm, but it allowed us to look for associations drawing on standard chemoinformatics-based searches. Surprisingly, of the 5,374 unique ingredients identified, only 2,966 were annotated as a primary suspect in at least one report; said a different way, 2,408 active drug ingredients had no reported ADRs whatsoever. A plot of the ingredients that were associated with ADRs shows that an exponentially decaying distribution, with 90% of the ADRs attributed to 40% of the drug ingredients (Supplementary Figure 1). After correction of distribution for ADRs with q-values better than 0.05, 90% may be attributed to 46% of the investigated drugs. This ingredient mapping was used throughout subsequent analyses (see Methods and Supplementary Material).

As expected, mapping drugs to their active ingredients, and not simply relying on synonym aggregation, reinforced the strength of the drug-ADR signals. For example, the non-steroidal anti-inflammatory drug (NSAID) indomethacin is used to treat chronic pain and fever^23^. When we assessed indomethacin as an ingredient, a strong signal linked it with gastric ulcer (RRR = 10.40; q-value = 3.65·10^-72^), and gastric ulcer hemorrhage (RRR = 7.99; q-value = 6.78·10^-18^). These adverse events are known from the labels of indomethacin-containing drugs, also confirmed in WDI^24^. However, when we searched the trade names of the drugs in which indomethacin is used (R_x_Norm synonym matching), these signals were dissipated in the noise: the strongest signal for gastric ulcer decreased to RRR = 1.79, q-value = 1.00; the strongest signal for gastric ulcer hemorrhage dropped to RRR = 2.42, q-value = 1.00.

### Bias in ADR reporting by indication, changes in regulatory, clinical, social and legal environment

Sometimes, ADRs are conflated with indications, and vice versa. An example is a report of rosiglitazone being prescribed for type 2 diabetes mellitus, with the ADR in the report being also diabetes mellitus (Table 1). Conversely, in another report rosiglitazone was identified as the primary suspect for congestive heart failure, as well as a therapeutic agent that was prescribed for the very same condition (Table 1). We quantified this indication bias both globally and over time. Approximately 5% of all reports for any drug describe the drug’s indication as an adverse event. The number of reports in which the same ADR and indication was reported increased linearly with the increasing number of yearly reports until 2011, followed by a sudden drop (Supplementary Figure 2). We took a closer look at the reports of rosiglitazone, where occurrence of diabetes as a side effect was attributed to the usage of this drug relatively frequently until 2004 (this obviously erroneous association is significant if considered in the reporting window of rosiglitazone (until 2011), with an RRR = 1.57 and a q-value < 10^-5^). After 2004 this association decreased, as did the overall prescriptions and reporting of this drug, owing to its widely-reported cardiovascular side effects^25,26^. In general, a simple comparison of indications and reported ADRs reduces the bias of verbatim repetition.

**Table 1.** Confusion of ADRs with indications. Report and case numbers identify two FAERS reports where the ADR is confused with the indication. For the first case, rosiglitazone prescribed for diabetes (Indication) is identified as the primary suspect (PS) for causing diabetes mellitus as an ADR as well. In the second case “cardiac failure congestive” is given as the indication for rosiglitazone with the reported ADR of “cardiac failure congestive”. The third case exemplifies correct reporting, where both the ADR and the indication of rosiglitazone are reported correctly.

| Report | Case | ADR | Drug / Role / Indication |
| --- | --- | --- | --- |
| 6545021 | 179039 | Diabetes mellitus | Rosiglitazone / PS / Diabetes mellitus |
| 5521616 | 162007 | Cardiac failure congestive | Rosiglitazone / PS / Cardiac failure congestive |
| 6380841 | 7085373 | Cardiac failure congestive | Rosiglitazone / PS / Diabetes mellitus |

We applied these methods to investigate how reports for individual drugs change over time. In particular, we monitored the total number of reports filed, and the incidence of adverse events preferentially reported at different time points. When reports are sorted by event dates in FAERS, “spikes” occur on the first day of each month, and even larger spikes on the first day of each year. Importantly, drug – serious ADR signals show a time-dependent increase (see Figures 2A, 3A and 4A, and 5A). The changes in drug-ADR associations over time can, of course, reflect new populations to which the drug is exposed.

We assessed the time evolution of reports of rofecoxib, a nonsteroidal anti-inflammatory drug (NSAID) that relieves pain through COX-2 inhibition (Figure 2). Several important events occurred over the clinical life of rofecoxib since its approval by the FDA in 1999: (1) A clinical study by Bombardier et al. published in November 2000 concluded that rofecoxib increased the risk of cardiovascular events^27^. (2) Introduction of warnings for cardiovascular events on the labels of Vioxx (a brand name of rofecoxib) in April 2002. (3) Withdrawal of rofecoxib from the market on September 30^th^ 2004.

**Figure 2.**
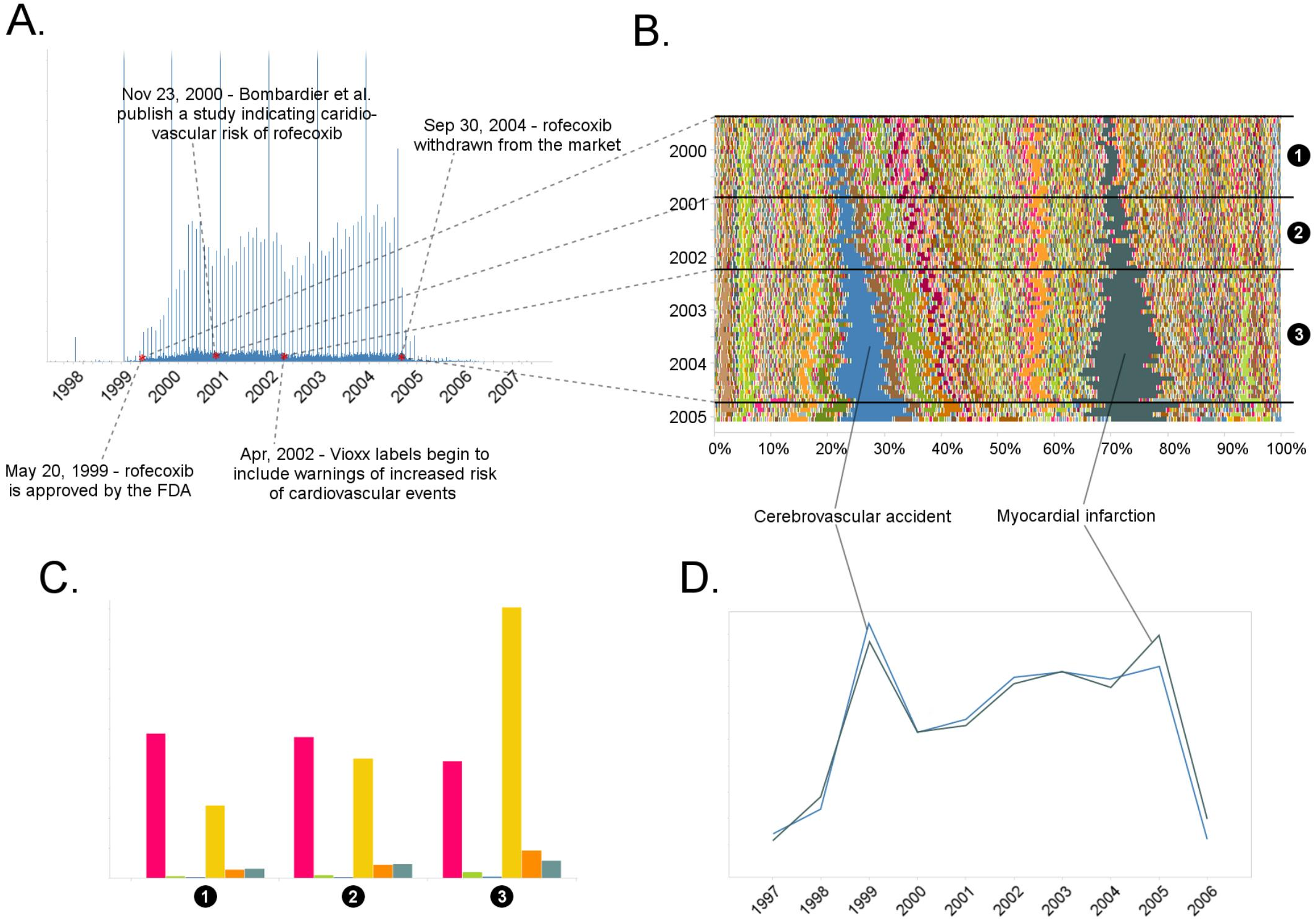
Submission pattern and time evolution of rofecoxib FAERS reports. A) Number of reports (per day) where rofecoxib was reported as primary suspect. Red dots represent events with a major impact on the FAERS reporting pattern of rofecoxib. B) Relative percent participation of all “preferred term” (PT)-level ADRs observed for rofecoxib. Each ADR is represented by a separate color. Characteristic time periods on the timeline of this drug are demarked by lines (associated with definitive events), and numbered. C) Identities of those reporting rofecoxib ADRs at the various reporting periods, marked to correspond with the annotations on panel B. Reports filed by physicians are shown in red; by pharmacists in blue; by other health-professionals in green; by lawyers in yellow; by customers in orange; reports in which the identity of the reporter was not stated are shown in dark blue. D) Enrichment-based clusters of ADRs (cerebrovascular accident and myocardial infarction) observed in rofecoxib reports between 1997 and 2006.

Myocardial infarction (RRR = 17.85; q-value < 10^-5^) and cerebrovascular accident (RRR = 17.69; q-value < 10^-5^) accounted for a large proportion of the ADRs reported for rofecoxib from its introduction in 1999 (Figure 2B and D). Before the study by Bombardier et al.^27^, most reports were filed by physicians. Between the Bombardier publication and the introduction of the label warning, these physician reports remained constant, while the number of reports by lawyers grew substantially. After the introduction of the label warning, the number of reports from physicians slightly decreased, but the trend to attribute myocardial infarction and cerebrovascular accident to administration of rofecoxib was further cemented by submitters who identified themselves as lawyers (see Figure 2C).

We also inspected the time evolution of another COX-2 inhibiting NSAID, celecoxib, approved by the FDA in December 1998, just shortly before Vioxx (Figure 3). Inspection of the timeline of celecoxib reports shows a slight increase in the number of reports around September 2004, reflecting the increase in use associated with the withdrawal of rofecoxib (Figure 3A). Until December 2004, the pattern of ADR in celecoxib reports is dominated by cerebrovascular accident (per-month RRR up to ~35) and myocardial infarction (per-month RRR up to ~45) in a similar fashion as in rofecoxib reports (Figure 3B and 3D). The increase of the overall number of reports around September 2004 coincided with concerns about the safety of celecoxib, likely reflecting a report of increased risk of cardiovascular events in patients who used celecoxib systematically over prolonged periods of time^28^. We checked whether the trends in reporting of side effects of celecoxib was affected by co-administration of rofecoxib, but the distribution of ADRs was almost identical after excluding the 8% of reports in which rofecoxib was present as a concomitant drug (Figure 3C). Closer examination of this pattern revealed that the reports during this period of time were largely stimulated by lawyers and “unidentified” individuals, while the contribution of health professionals remained steady much below the level of reports for rofecoxib (Figure 3E). These trends were confirmed by logistic regression modeling (see Methods and Supplementary Table 1), which showed that reports of myocardial infarction were significantly correlated with reports of celecoxib filed by lawyers before 2005 (Model 4 in Supplementary Table 1).

**Figure 3.**
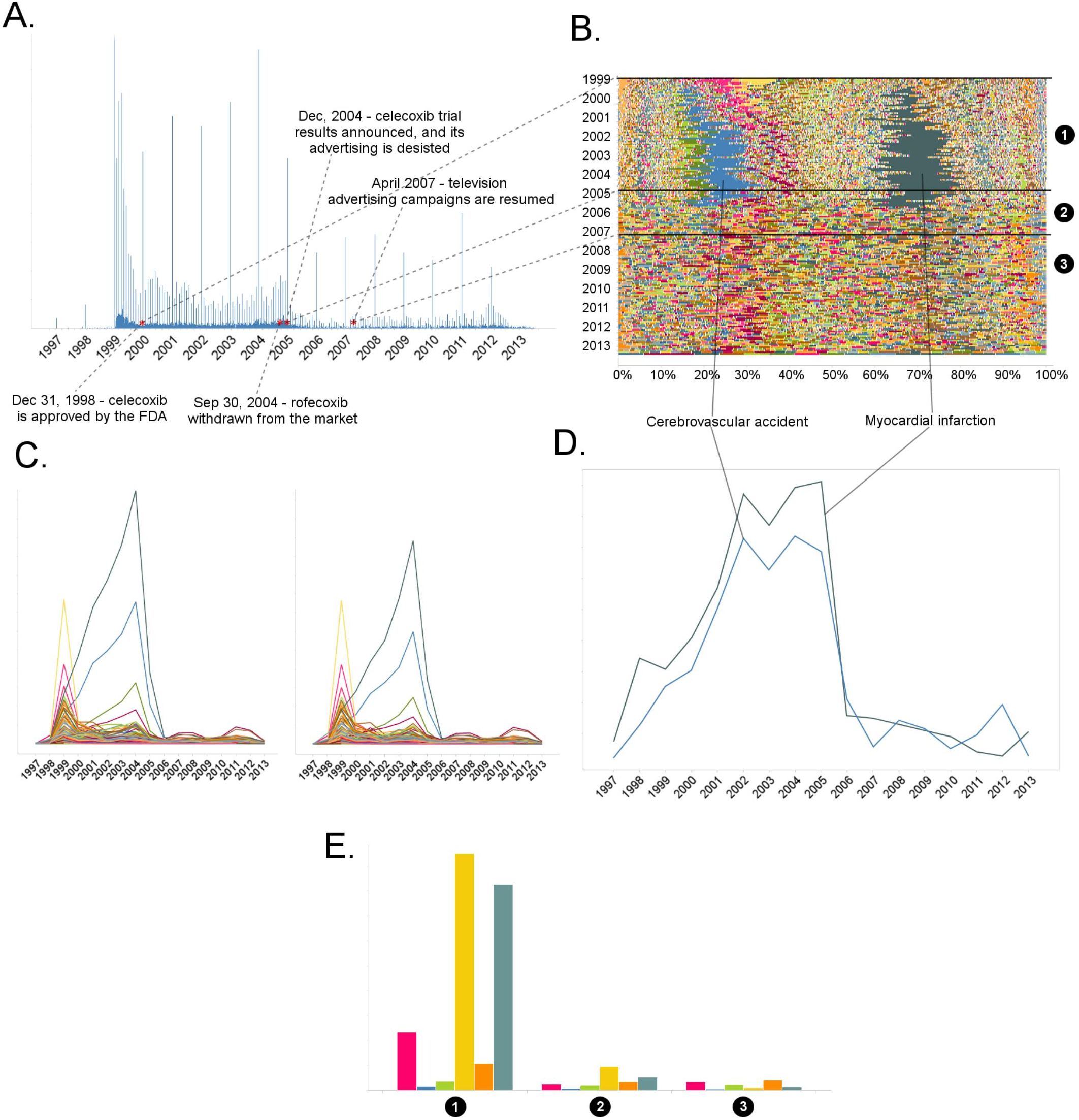
History of FAERS reports on celecoxib. A) Number of FAERS reports (per day) where celecoxib was reported as primary suspect. B) Relative percent participation of all PT-level ADRs observed for celecoxib. Each ADR is represented by a separate color. Characteristic time periods on the timeline of this drug are marked by lines, and numbered. C) Per-month number of reports where celecoxib was primary suspect; each line corresponds to a separate PT-level ADR. The left-hand plot describes all reports with celecoxib as primary suspect. In the plot on the right the reports in which rofecoxib was also present were omitted. Colors are matched with those used in panel B. D) Enrichment-based clusters of most frequently reported ADRs (cerebrovascular accident and myocardial infarction) observed in rofecoxib reports. Colors match those in B and C. E) Identities of those reporting celecoxib ADRs at various reporting periods, marked to correspond with the annotations in panel B. Reports filed by physicians are shown in red; by pharmacists in blue; by other health-professionals in green; by lawyers in yellow; by customers in orange; reports in which the identity of the reporter was not stated are shown in dark blue. The distributions are shown for various time periods, annotated with similar numbers in black circles in panel B.

### Drugs with similar chemical structure and modes of action may display distinct clinical ADR phenotypes

It is generally expected that compounds with similar structures and modes of action will have similar ADR profiles; for instance, several selective serotonin reuptake inhibitors (SSRIs) are associated with suicidal behavior in young adults^29,30^. However, this is not always the case. The post-marketing ADR reports of the structural analogs rosiglitazone^31^ and pioglitazone, which act on the same primary target (PPAR-γ) and are structurally related, are notably different (compare Figure 4B and 5B).

**Figure 4.**
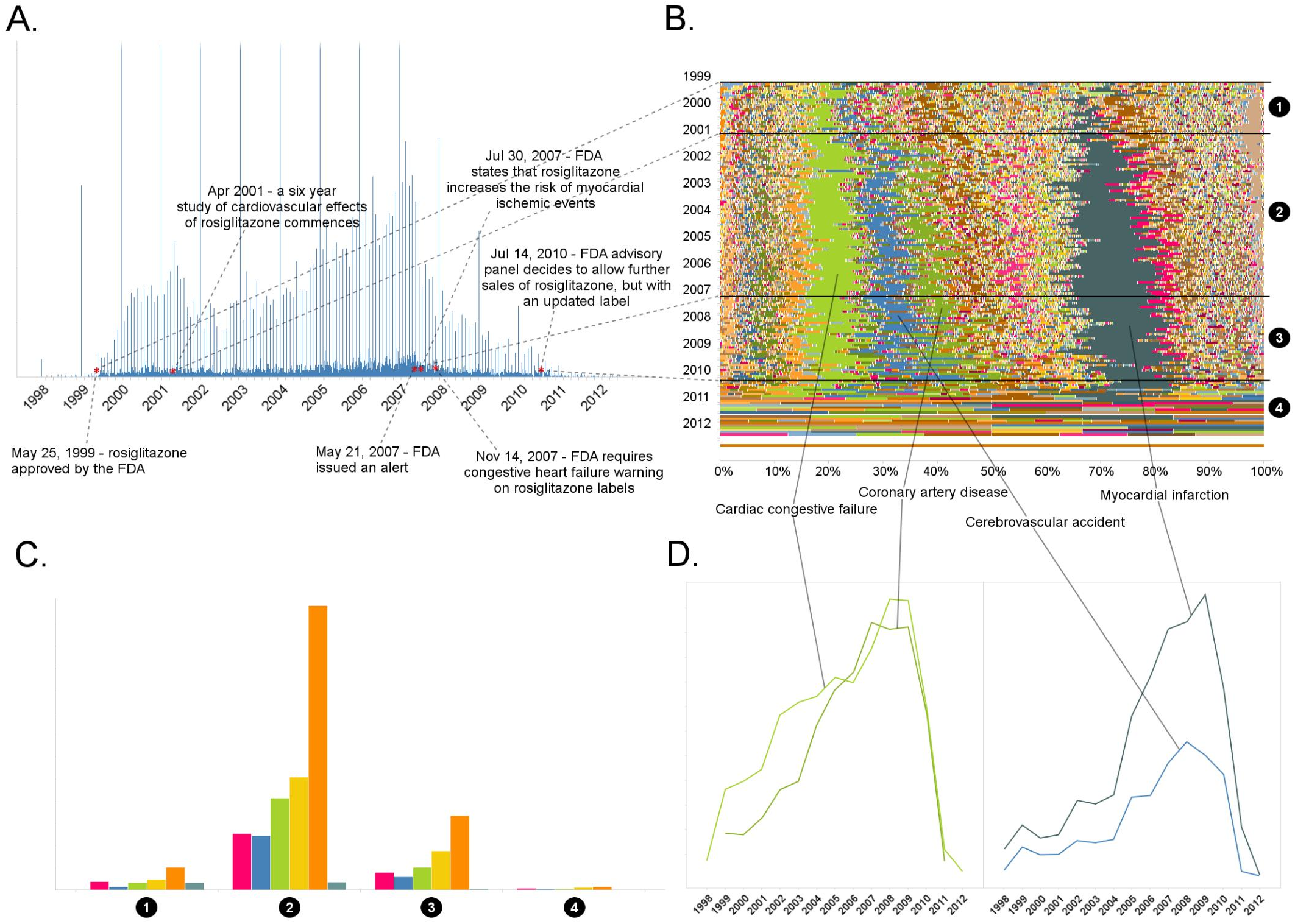
Rosiglitazone reports. A) Number of FAERS reports (per day) where rosiglitazone was reported as primary suspect. B) Per-month percent participation of all PT-level ADRs observed for rosiglitazone. Each ADR is represented by a separate color. Characteristic time periods on the timeline of this drug are demarked by lines, and numbered. C) Identities of those reporting rosiglitazone ADRs at various reporting periods. Reports filed by physicians are shown in red; by pharmacists in blue; by other health-professionals in green; by lawyers in yellow; by customers in orange; reports in which the identity of the reporter was not stated are shown in dark blue. The distributions are shown for various time periods, and are annotated with numbers in black circles that correspond to time periods annotated with similar numbers in black circles in panel B. D) Enrichment-based clusters of ADRs observed in rosiglitazone reports.

For rosiglitazone, many heart-related reports have been filed since its FDA approval in May 1999 (Figure 4A, 4B). Whereas the absolute number of reports have varied over time, and has been affected by the clinical trial and scientific reports in much the same way as rofecoxib, the predominance of heart effects, such as congestive cardiac failure (RRR = 31.99; q-value < 10^-5^), coronary artery disease (RRR = 26.32; q-value < 10^-5^), cerebrovascular accident (RRR = 11.72; q-value < 10^-5^), and myocardial infarction (RRR = 20.73; q-value < 10^-5^), relative to other events, has been unperturbed throughout the lifetime of this drug (Figure 4B and D).

The other hypoglycemic drug, pioglitazone, has triggered fewer reports of heart effects relative to the clinical ADR profile of rosiglitazone since its approval in July 1999 (Figure 5A). Whereas analysis of FAERS reports does support a statistically significant signal between pioglitazone and cardiac failure (RRR = 5.09; q-value < 10^-5^), the time evolution of this signal reveals that the major contribution to its statistical strength comes from a single peak that subsides by the year 2002, and coincides with the increased scrutiny of rosiglitazone (Figure 5C). Unlike rosiglitazone, the ADR landscape of pioglitazone is dominated by bladder cancer (RRR = 305.69; q-value < 10^-5^), with a substantial increase in reports from 2009 onward (Figure 5B). Conversely, this signal is significantly underrepresented in the rosiglitazone reports (RRR = 0.12; q-value < 10^-5^; see Figure B). There is evidence that non-selective PPAR agonists (α + γ) such as pioglitazone could contribute to carcinogenesis^32^, and a recent study linked bladder cancer to the development of chronic kidney disease as an effect of long term use of pioglitazone^33^. Still, the mechanisms linking the less selective pioglitazone but not the selective PPAR-γ agonist rosiglitazone to bladder cancer are unclear, and this association must remain tentative.

**Figure 5.**
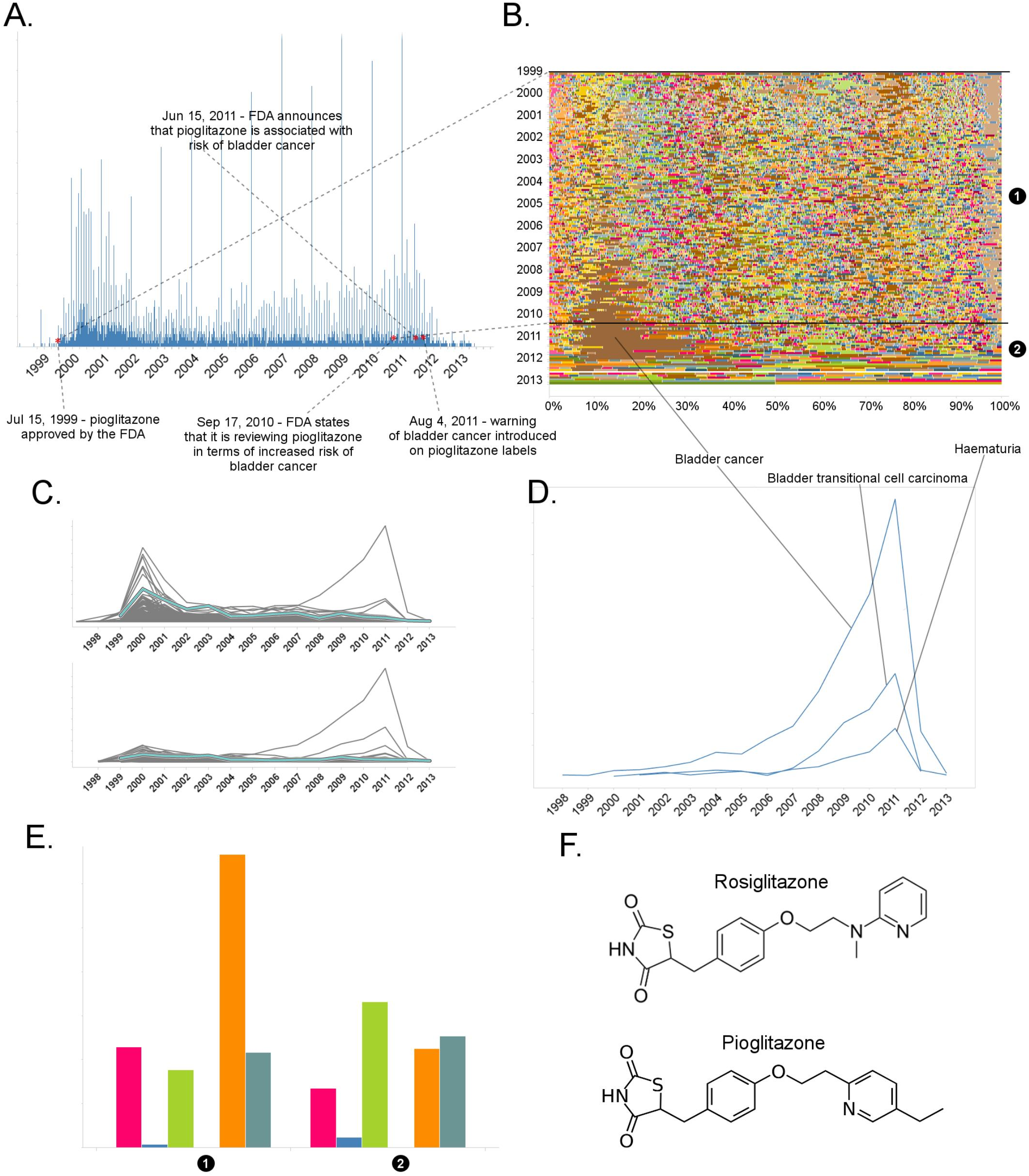
The landscape of pioglitazone reports. A) Number of FAERS reports (per day) where pioglitazone was reported as primary suspect. B) Per-month percent participation of all PT-level ADRs observed for pioglitazone. Each ADR is represented by a separate color. Characteristic time periods on the timeline of this drug are marked by lines, and numbered. C) Per-month number of reports where pioglitazone was primary suspect; each line corresponds to a separate PT-level ADR. The plot on the top of the panel shows number of times individual ADRs have been reported, and the bottom the corresponding per-month enrichments. The traces for cardiac failure have been distinguished by the blue color. D) Enrichment-based clusters of cancer-related ADRs observed in pioglitazone reports. E) Identities of those reporting pioglitazone ADRs at various reporting periods. Reports filed by physicians are shown in red; by pharmacists in blue; by other health-professionals in green; by lawyers in yellow; by customers in orange; reports in which the identity of the reporter was not stated are shown in dark blue. The distributions are shown for various time periods, and are annotated with numbers in black circles that correspond to time periods annotated with similar numbers in black circles in panel B. F) Structure of rosiglitazone and pioglitazone.

### Using monthly report counts to de-bias stimulated reporting

The trends and biases in ADR reporting can hamper the division and reliability of drug-ADR associations. The statistically significant association that we found between pioglitazone and cardiac failure stems mostly from the reports from before 2004, which can be attributed to the popularity of the contention that hypoglycemic fibrates cause cardiovascular side effects^26^. We calculated month-resolved statistical significance of the pioglitazone – cardiac failure association. The majority of considered dates indicated that there was no statistically significant association between this drug-ADR pair (top panel in Supplementary Figure 3). Conversely, the association of rosiglitazone and myocardial infarction was statistically significant in nearly every time period (bottom panel in Supplementary Figure 3). The periods where the pioglita-zone - cardiac failure association is statistically significant are contained to a couple of sparse spikes (Supplementary Figure 3), and so we consider this association to be stimulated, and essentially artifactual. Such month-resolved statistical analysis for drug-ADR associations may be broadly helpful in detecting biased reporting trends.

### Combining pharmacokinetics & FAERS to investigate mechanism and for reverse translation

There is great interest in using pharmacovigilance for target identification and to illuminate therapeutic and ADR mechanism of action^30,34–37^. By matching to *in vitro* activity, one may hope to link an ADR that emerges in FAERS with the targets responsible for the physiology, making the linkage: drug→ known target→ ADR. Whereas we ourselves have championed the role of *in vitro* pharmacology for anticipating possible toxicology^38–40^, doing this reliably depends on knowing the exposure of the drug to the implicated target. Without considering drug pharmacokinetics, FAERS-based inference of drug→ target→ ADR associations can mislead^41^.

An illustrative example is the hypertension associated with inhibition of the VEGF-R2 (see Methods). The relevance of such inhibition to hypertension is supported by the high-incidence of this ADR with the VEGF-R2 specific antibody, bevacizumab (Table 2), and is well-accepted^42^. Correspondingly, some small molecule kinase inhibitors that inhibit VEGF-R2 with relevant in vivo pharmacokinetics, such as Pazopanib and Sorafenib and Sunitinib (Table 2), also share a hypertension ADR. However, other kinase inhibitors with VEGF-R2 inhibition do not appear to increase the reported incidence of hypertension (Table 2). High incidence is reported only with those drugs in this class which have exposure margins (EM) less than 13 for this target (biochemical IC_50_/C_max_; Table 2). Using such an EM cutoff in the FAERS analysis, the signal for this ADR over random will separate drugs with true adverse event from those which lack it (Table 2).

**Table 2.** Hypertension associated with VEGF-R2 inhibition depends on the exposure margin of small molecule anti-VEGF-R2 drugs (VEGF-R2 IC_50_/C_max_). FAERS reports of small molecule kinase inhibitors with VEGF-R2 inhibition show an increased incidence of hypertension reports only in case their exposure margin is less than 13. The label of drugs with high incidence of hypertension in FAERS lists this side effect, while none of those drugs which have low incidence carry the label. *p-value of association between drug and hypertension < 0.001. Counts (N), expected counts (E), and an often-used disproportionality measure (EB05) based on the FDA’s FAERS database of spontaneous reports of suspected drug adverse drug reactions are provided. The values of E are the expected number of patients reporting vascular hypertensive disorder after taking each drug if the drug reports and the reports of the event were independent within the database, conditional on the patients age and gender. The ratio N/E is a measure of disproportionality of report counts of each particular drug-event combination. The value EB05 (empirical Bayes 5% lower bound of a 90% credible interval) is a conservative estimate of the true reporting disproportionality that uses estimated overall prevalence of drug-ADR associations throughout the database. The value of EB05 is less than N/E and has the effect of correcting the simple ratio for sampling variance and multiple comparisons bias. See ^43–46^ for details and discussion of the FAERS database and the use of disproportionality analyses within spontaneous report databases. The values of EB05 for the first three drugs indicate 95% confidence that reports of those three drug-event combinations are reported about 3 or 4 times as often as would be expected if they were independent, while the values of EB05 < 1 in the final three drugs in the table indicate no evidence for higher than expected reporting rates. ^†^Significant increase.

| Drug | VEGF-R2 IC<sub>50</sub><br>(μM) | Free C<sub>max</sub><br>(μM) | Exposure Margin<br>(EM) | N | E | EB05 | FAERS data | | |
| --- | --- | --- | --- | --- | --- | --- | --- | --- | --- |
|  |  |  |  |  |  |  | #reports of hypertension | Total number of reports | %hypertension of total reports |
| Pazopanib | 0.0069 | 0.13 | 0.053 | 576 | 123 | 4.329<sup>†</sup> | 564 | 10534 | 5.35* |
| Sorafenib | 0.025 | 0.055 | 0.45 | 794 | 183.48 | 4.055<sup>†</sup> | 700 | 10599 | 6.60* |
| Sunitinib | 0.015 | 0.006 | 2.5 | 1000 | 305.84 | 3.093<sup>†</sup> | 914 | 21420 | 4.27* |
| Gefitinib | 2.5 | 0.19 | 13 | 25 | 82.153 | 0.227 | 14 | 3998 | 0.35 |
| Crizotinib | 1.4 | 0.08 | 18 | 10 | 50.874 | 0.131 | 9 | 4138 | 0.22 |
| Dasatinib | 1.7 | 0.006 | 283 | 68 | 115.5 | 0.485 | 54 | 8583 | 0.63 |

A more complex case emerges through the investigation of methylphenidate and the atypical antipsychotics, risperidone/paliperidone^47^ and aripiprazole, drugs prescribed for the treatment of attention deficit hyperactivity disorder (ADHD)^48–50^. FAERS analysis indicates that treatment with risperidone/paliperidone, the latter of which is the main active metabolite of risperidone, increases the frequency of gynecomastia and galactorrhea, while methylphenidate has a low incidence of these ADRs, as do other atypical antipsychotics, such as aripiprazole (Figure 6)^9^). For example, between 2007 to 2013 there were respectively 5,073 and 123 cases in FAERS where risperidone and paliperidone are the primary suspect of gynecomastia (Figure 6, RRR = 113.82, q-value < 10^-5^; RRR = 7.53, q-value <10^-5^). For aripiprazole (RRR = 0.85) and methylphenidate (RRR = 1.39), however, the q-values were close to 1, indicating no significant associations with this ADR (Figure 6B). Thus, the FAERS data clearly separates the profile of risperidone/paliperidone from both methylphenidate, with which it overlaps for treatment of ADHD, and from other atypical antipsychotics, like aripiprazole. The inference would be that the target responsible for the gynecomastia and galactorrhea for risperidone/paliperidone is not modulated by either methylphenidate or any other atypical antipsychotics. Whereas this is correct for methylphenidate, it is incorrect for the atypical antipsychotics.

**Figure 6.**
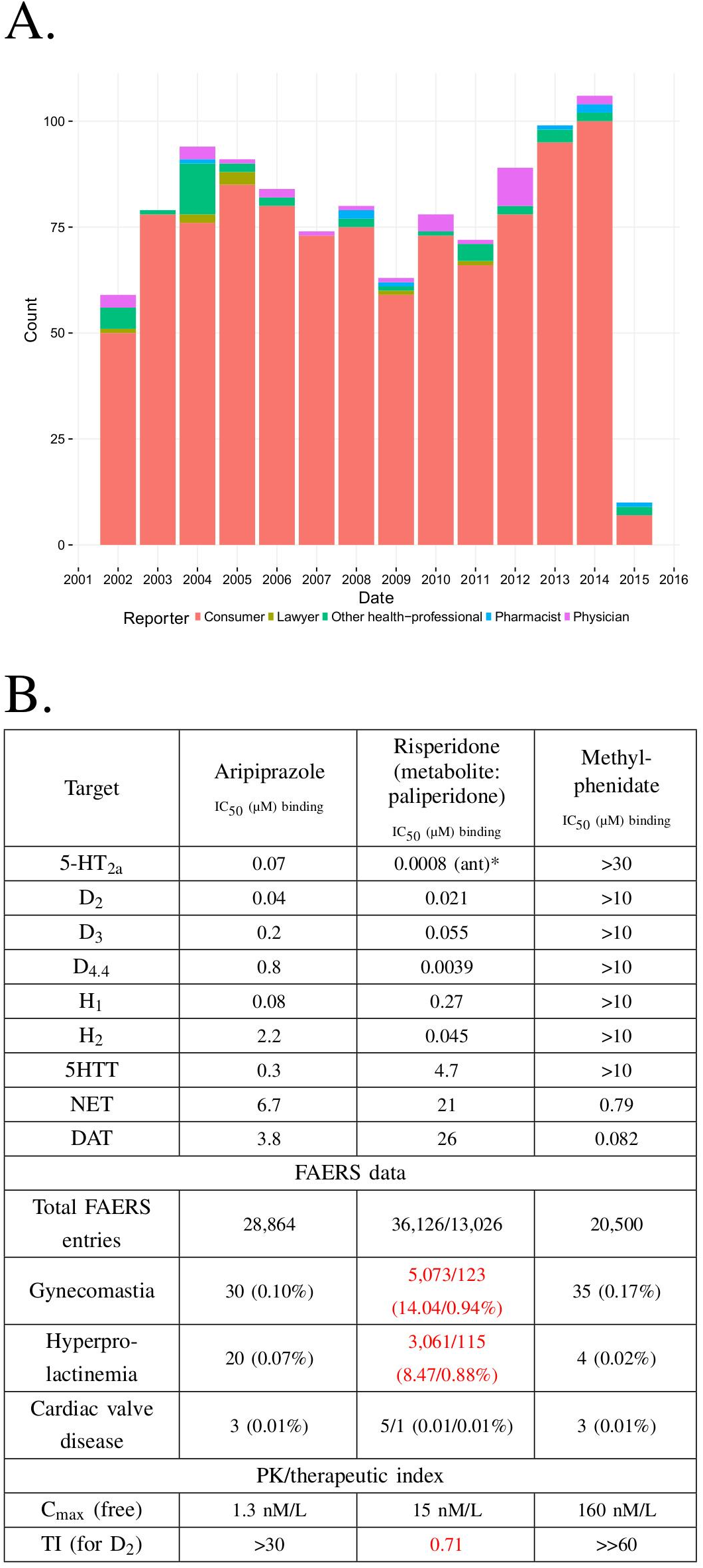
Integration of pharmacodynamic and pharmacokinetic data is necessary to interpret FAERS information. A) FAERS analysis of the reporting pattern of gynecomastia in patients treated with risperidone between 2002-2015. B) Summary table of the *in vitro* pharmacological profile, FAERS entries (total number of reports, and reports of gynecomastia, hyperprolactinemia and cardiac valve disease where the listed drugs were the primary suspects) and calculation of therapeutic index of aripiprazole, risperidone/paliperidone and methylphenidate. The prominent effects of risperidone/paliperidone at the D_2_ dopamine receptor in conjunction of the narrow TI differentiates these compound(s) from the rest. Assays were performed at the Novartis Institutes for BioMedical Research, Cambridge. *Asterisks denote functional assays. ant: antagonism; ago: agonism

Both aripiprazole and risperidone/paliperidone are atypical antipsychotics with high affinity to dopaminergic, serotoner-gic, adrenergic and histaminergic receptors (Figure 6B^38, 51^). It is well known that inhibition of the D_2_ dopamine receptor is linked to hyperprolactinemia, which is the underlying mechanism of gynecomastia and galactorrhea^52^. Regardless of their similar potency at the D_2_ dopamine receptor the difference between the ADR profile of aripiprazole and risperidone/paliperidone is explained by their mechanism of action (risperidone is a full antagonist and aripiprazole is a partial agonist) and particularly by their PK profile, which reveals that the exposure margin (EM^41^) for D_2_ is large for aripiprazole and so this ADR did not manifest. For risperidone, the TI is less than 1 which explains the high incidence of gynecomastia (Figure 6). Methylphenidate does not affect the D_2_ receptor at all, thus this ADR was not observed.

## DISCUSSION

Four key observations emerge from this study. First, much of the potential signal in FAERS and related databases is obscured by chemical name redundancy. This introduces false associations that would fall to insignificance on synonym aggregation, and hides associations that would be significant on aggregation. This may be addressed by representing active ingredients by their unique chemical structures in a readily searchable form (SI File 1). Second, FAERS reports tilt toward serious outcomes, partly owing to a confusion of ADRs and outcomes. Third, FAERS suffers from several forms of conflation: multiple entries, indications with ADRs, newsworthiness, and scientific and legal influences. These may be detected by statistical analyses, including comparing reports over time. Fourth, and perhaps more generatively, once these biases and conflations are corrected, the molecular mechanism of previously hidden ADRs can be revealed; an example explored here is the association of urinary bladder cancer with mixed PPAR-α and PPAR-γ agonists.

A major reason for FAERS’s bias toward serious outcomes is the conflation of ADRs and outcomes. This may stem from an issue as simple as confusion on whether “death” is listed as an ADR – associated with the drug only – or an outcome – associated with the disease itself. This is the case with the attribution of the side effect “death” to thalidomide’s use in complex myeloma multiplex, when this reflects the high mortality rate of the disease itself^31^. Naturally, there are some cases where use of a drug can increase death rate, even in treating life-threatening diseases, such as the case of milrinone for acute heart failure syndromes (AHFS)^53^ or severe chronic heart failure^54^. Accordingly, the category “outcome” should be used cautiously for ADR analysis, especially in large-scale studies that aggregate data from several drugs.

In principle, submission of FAERS reports requires medical knowledge, as they include specific indications for which drugs were prescribed, identification of the primary suspect of ADRs, and structured description of ADRs by MedDRA terms. Nevertheless, a third of the reports are contributed by customers, and a half by submitters who do not identify themselves as medical professionals, including lawyers. This contributes to the high redundancy and error in FAERS, and to the “stimulated reporting” from which it suffers^55^. This appears to have been the case with celecoxib, whose association with cerebro− and cardiovascular events in FAERS reports was driven primarily by reports from legal professionals (Figure 3E). After rofecoxib was withdrawn, the proportion of these events for celecoxib returned to background. For cases like these, a temporal analysis of ADR-drug associations can illuminate spurious associations. Interrogation of FAERS and related databases to illuminate the molecular mechanisms of ADRs, and indeed the shared target profiles of drugs, has been an area of much recent interest^56^. Here, too, we find that the disambiguation of ADRs, indications, reporting and indication biases can reveal previously obscured associations. An example is the association of bladder cancer with the mixed PPAR-α and PPAR-γ agonist pioglitazone. FAERS analysis is instrumental here, providing information on a large patient population and enabling the comparison with the selective PPAR-γ agonist rosiglitazone, which is not associated with bladder cancer (Figure 5F)^57^.

Improvement of statistical methods for signal detection is an area of active research^2–4,6,58^ and a lot of attention is payed to advanced statistical methods such as (Bayesian) information components^58^, Empirical Bayes statistics^59^, and hierarchical methods^59^. As with all machine learning and statistical approaches, these methods assume clean input data – the biases and noise they address is of statistical nature. We have used a well-known disproportionality approach, relative reporting ratio (RRR) with 
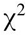
 test statistic for disproportionality. The RRR has its limitations and may underperform compared to more advanced methods^58^. The focus of our study was how proper preparation of the input data – cleaning drug ingredient mapping, and estimating multiple reporting – boosts signal detection performance, even with a simple method such as the RRR. We believe that applying the procedures and precautions we described here together with more advanced statistical methods will boost their performance even further.

A key caution is that to be relevant, not only must one associate ADRs and drugs, and drugs and their targets, but one must ensure that the pharmacokinetics of the drugs ensure that the implicated target is exposed to the drug at relevant concentrations. This is illustrated by the comparison of the ADHD drugs risperidone and aripiprazole and gynecomastia. Both drugs affect the D_2_ dopamine receptor that underlies the ADR, but only risperidone reaches a sufficient exposure to trigger it. The VEGF-R2 inhibitor example confirms that this type of evaluation is the only way one can objectively detect ADR-target pairs and explain the underlying mechanisms of their manifestation. Relying only on a ADR→ drug→ *in vitro* target schema can be insufficient to understand shared targets or molecular mechanisms; as Goodman long ago suggested, pharmacokinetic exposure remains crucial^60^.

## CONCLUSIONS

The challenges and opportunities in FAERS and indeed from related databases, flow from its ambitions. It publishes multiple reports - physicians, patients, other medical professionals, attorneys - on multiple drugs, named in multiple ways and taken in multiple contexts. FAERS does not represent a strictly reviewed and carefully channeled source of observations about drugs, as a clinical trial does - there is no placebo arm in FAERS, nor are there reports of cases when a given drug was prescribed and caused no side effects. It contains uncontrolled, volunteered information on a large scale. This may be seen as a feature of FAERS - a database designed with hypothesis generation rather than hypothesis testing in mind. Still, the hypotheses that FAERS suggests depend critically on the ability to disentangle its data. Tools like those described here are crucial to control for the often conflated and contradictory observations in FAERS reports, where serious outcomes are over-reported, reported death is often linked to submission by the patients themselves, a single event is reported multiple times, true associations between drugs and adverse events are missed because a single agent is named in multiple ways, or a mechanistically related disease occurs in different system organ categories. Once its data are disentangled, FAERS represents unprecedented opportunities to track drug outcomes in large patient populations, revealing new associations. The power of such analysis is that it may be applied systematically and comprehensively across a massive number of observations.

We recognize that a fully automated method, such as that described here, cannot replace expert knowledge. What such a method can do is identify, prioritize and sometimes deprioritize drug-adverse event associations, and sometimes even mechanistic inference, for detailed expert identification. This approach should be useful to the growing community of regulators, payers, physicians, and patients that work with and depend upon trends emerging in FAERS to improve drug use and health outcomes. By making several of these tools available to the community, we hope to enable future interrogation of FAERS by other investigators.

## METHODS

### FAERS data source

FAERS reports were downloaded on May 24^th^ 2016 from the FEARS database ^8^ for the years between 4^th^ quarter of 1997 and 4^th^ quarter of 2015, inclusive. ADR, indication, drug role (primary suspect, secondary suspect, concomitant), and outcome data was mapped using ISR report identifiers to the individual reports. Drugs were identified by the reported drug name in FAERS.

### Mapping drugs to ingredients

We assembled a list of synonyms of drugs, using public and licensed databases including Thompson Reuters Integrity ^61^, GVK ^62^, Drugbank ^63^, ChEMBL ^64^, and R_x_Norm^22^. These synonyms were matched with drug products, and constituent molecular ingredient structures, encoded using as InChIKeys ^65^. We read in all the drug names from all the FAERS reports, and all the synonyms that had been assembled. Non-alphabetical characters (except numbers), capitalization, and terms that carried little information regarding the identity of the drugs (such as articles, or often occurring words like “acid”) were removed from the FAERS drug names and synonym names, and the remaining parts of the names were tokenized. Each tokenized FAERS drug was then compared to each tokenized synonym, and the overlap of tokens was recorded for each pair using the Tanimoto similarity coefficient *t_c_*. The synonym with the highest *t_c_* value was picked for a given drug, as long as the *t_c_* was ≥0.2; for any drug, if a synonym with *t_c_* of 0.99 or higher was found, it was considered to be an exact match, and used to identify the drug in question without comparison to further synonyms. For the most frequent (among the top 500) drug names in FAERS, we manually mapped those drug names to InChIKeys that could not be mapped. Since InChIKeys are not typically calculated for large macromolecules, we used the non-proprietary name in lieu of the InChIKey in these cases.

### Adverse drug reaction terms

The majority of ADRs in FAERS are reported using the Medical Dictionary for Regulatory Activities (MedDRA) ^66^. Some older reports contained terms that are not part of the newer MedDRA that is used currently. To normalize and annotate the ADR terms extracted from reports we used a Levenshtein algorithm that compared the FAERS ADR terms to the MedDRA terminology. We set the minimal Levenshtein score at which a given MedDRA term was considered a perfect match to 0.95, and the minimal acceptable score to 0.90 at above which the highest scoring term was picked to standardize a given ADR. Additional 32 ADR terms were standardized manually, leaving less than 0.5% ADR terms unmatched.

### Establishing ingredient - ADR and ingredient - indication associations

We used the well-established Relative reporting ratio (RRR) together with a 
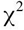
 statistic for disproportionality signal detection ^58^. We constructed ingredient-ADR contingency tables and calculated the expected number of occurrences, the RRR, and Yates-corrected 
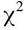
 p-values ^67^ for these contingency tables, as implemented in SciPy ^68^. False discovery rate (FDR) was controlled ^68^ using the Holm procedure ^69^, yielding q-values. Associations were selected if they: a) were reported at least 5 times in FAERS; b) had a q-value < 0.05; and c) had an RRR > 1. These ingredient - ADR pairs are shown in Supplementary File 2.

### Calculating ingredient – ADR associations on monthly basis

With FAERS data annotated with dates of ADRs, for every ingredient – ADR pair we calculated co-occurrence frequencies, RRR-values, and 
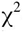
 based p-values for every month between January 1997, and December 2015. In these calculations we used the numbers of drugs, ADRs, and total reports from the relevant month only. False discovery rate (FDR) was controlled using the Holm procedure (for each month separately), yielding q-values. In Supplementary File 3 for every statistically significant (in aggregate) ingredient – ADR association we reported the numbers of months where q-values were lower than 0.05, and where q-values were higher than or equal to 0.05.

### Clustering of ADR by time evolution

We considered the numbers of reports in each month (time evolutions) for individual ADRs observed across FAERS for four drugs: rofecoxib, celecoxib, rosiglitazone, and pioglitazone. With knowledge of numbers of each of the considered drugs, ADRs, and the number of total reports in FAERS in each month, we calculated month-resolved RRR-values for the drug-ADR pairs. The time evolutions of the RRR-values were clustered for each drug using the partitioning method for maximum dissimilarity, as implemented in R ^70^ scored by the similarity (Pearson correlation coefficient) of time evolutions of RRR.

### Logistic regression models of myocardial infarction dependence on the use of celecoxib

For every FAERS report, we noted whether a) celecoxib was reported as the primary suspect drug, b) whether myocardial infarction was reported, c) the occupation of the person filing the report, and d) whether the reported event took place before 2005 (when rofecoxib was still on the market). Using this data and R’s implementation of binomial logistic regression (via the glm() function), we prepared four models (with the logit link function) ^70^ to investigate if myocardial infarction is associated with the use of celecoxib, the occupation of the person filing the report, and the report being filed before 2005. In each model, reporting myocardial infarction served as the output variable, and combinations of the remaining variables were used as input variables. Resulting models are summarized and described in more detail in Supplementary Table 1.

### Analysis of association between VEGF-R2 inhibition and hypertension

Apart from analysis described in this work, additional Bayesian data mining and statistical analysis of VEGF-R2 inhibition-related hypertension was based on the methods described in detail by DuMouchel ^43^, DuMouchel and Pregibon ^44^, Szarfman et al. ^45^, and Almenoff et al. ^46^

## ACKNOWLEDGEMENTS

AM was supported in the initial stage of this work by the NIBR Presidential Postdoctoral Fellowship co-mentored by LU and BKS. The work of BKS was supported by US National Institutes of Health Grant R01 GM71896. Authors thank Dr Duncan Armstrong for his comments and advice on the manuscript.

## CONFLICTS OF INTEREST

BKS has previously consulted for Novartis.

